# Antibiotics modulate attractive interactions in bacterial colonies affecting survivability under combined treatment

**DOI:** 10.1101/2020.12.14.422681

**Authors:** Tom Cronenberg, Marc Hennes, Isabelle Wielert, Berenike Maier

## Abstract

Biofilm formation protects bacteria from antibiotics. Very little is known about the response of biofilm-dwelling bacteria to antibiotics at the single cell level. Here, we developed a cell-tracking approach to investigate how antibiotics affect structure and dynamics of colonies formed by the human pathogen *Neisseria gonorrhoeae*. Antibiotics targeting different cellular functions enlarge the cell volumes and modulate within-colony motility. Focusing on azithromycin and ceftriaxone, we identify changes in type 4 pilus (T4P) mediated cell-to-cell attraction as the molecular mechanism for different effects on motility. By using strongly attractive mutant strains, we reveal that the survivability under ceftriaxone treatment depends on motility. Combining our results, we find that sequential treatment with azithromycin and ceftriaxone is synergistic. Taken together, we demonstrate that antibiotics modulate T4P-mediated attractions and hence cell motility and colony fluidity.

**Author Summary:** Aggregation into colonies and biofilms can enhance bacterial survivability under antibiotic treatment. Aggregation requires modulation of attractive forces between neighboring bacteria, yet the link between attraction and survivability is poorly characterized. Here, we quantify these attractive interactions and show that different antibiotics enhance or inhibit them. Live-cell tracking of single cells in spherical colonies enables us to correlate attractive interactions with single cell motility and colony fluidity. Even moderate changes in cell-to-cell attraction caused by antibiotics strongly impact on within-colony motility. Vice versa, we reveal that motility correlates with survivability under antibiotic treatment. In summary, we demonstrate a link between cellular attraction, colony fluidity, and survivability with the potential to optimize the treatment strategy of commonly used drug combinations.

## Introduction

Aggregation has a strong potential for protecting bacteria from antibiotics. A multitude of different mechanisms has been shown to be responsible for protection [1]. They include inhibition of penetration of antibiotics into the biofilm [2, 3] and activation of stress responses [4-6]. Reduction of growth rate, rate of protein synthesis, and proton motive force [1] often enhance tolerance against antibiotics [7, 8] and increase the survival time during antibiotic treatment.

Some antibiotics cause structural rearrangements in biofilms. In particular, antibiotic treatment can induce biofilm formation [9, 10]. Very little is known, however, about how antibiotics affect biofilm structure at the single cell level. Only recently it was shown that the local structure of *Vibrio cholerae* biofilms rearranges upon antibiotic treatment [11]. In particular, under treatment with proteins synthesis inhibitors the density of bacteria was reduced by a combined effect of increasing cell volume and decreasing amount of the matrix component that is responsible for cell-cell attraction in *V. cholerae* biofilms [11]. Lacking single cell resolution, this cell volume increase would have been observed as an increase in biofilm mass and likely interpreted as antibiotic-induced biofilm formation.

While first reports of biofilm structure at the single cell level are emerging, it is unclear how antibiotics affect biofilm dynamics. Many bacterial species use type 4 pili (T4P) for generating attractive forces between cells that initiate and stabilize biofilms [12]. Next to forming T4P-T4P bonds between neighboring cells, T4P function as strong molecular machines [13, 14] that govern the viscosity or fluidity of colonies [15-17], i.e. early biofilms. T4P polymers grow, bind, and retract continuously [18], enabling a tug-of-war between bacteria in a colony [14, 19]. For *Neisseria* species it was demonstrated that by modulating motor activity, T4P-T4P binding force, or T4P density, the dynamics of a colony transitions from gas-like to fluid-like to solid-like [15-17]. It is tempting to speculate that colony fluidity affects bacterial fitness. Indeed, it was shown that fluidity correlates with the ability to colonize narrow vessels [15]. It is also conceivable, that high motility of cells within colonies enables rearrangements of cells during growth and division and allows bacteria residing at the center of the colony to reproduce. Moreover, cell motility has been connected with enhanced diffusion of macromolecules into biofilms [20] supporting influx of both nutrients and antibiotics. Therefore, we address the question how bacterial motility affects survival under benign conditions and under antibiotic treatment.

*Neisseria gonorrhoeae* (gonococcus) is the causative agent of gonorrhea and conjunctivitis. Gonorrhea is currently the second most common sexually transmitted disease and the probability for failure of antibiotic treatment is rising rapidly [21]. Some aspects of antibiotic resistance are understood at the genetic level [22-25], but the effect of colony and biofilm formation, its structure, and its dynamics on the efficiency of treatment remain largely unexplored.

In this study, we employ a single cell approach to characterize effects of antibiotic treatment on colony structure and dynamics of *N. gonorrhoeae*. We show that antibiotics with different targets consistently affect cell volume and colony structure. Different antibiotics have varying effects on the cellular dynamics within the colonies. Focusing on the two currently recommended antibiotics for gonorrhea treatment, azithromycin and ceftriaxone [26], we correlate dynamics to T4P-T4P interactions by showing that antibiotic treatment can reduce or enhance T4P-mediated cell-to-cell attraction. Cellular dynamics itself impacts on the efficacy of antibiotic treatment. Motivated by the motility-enhancing effect of azithromycin, we study sequential treatment with azithromycin and ceftriaxone and find synergistic effects.

## Results

### Antibiotic treatment affects biofilm structure and cell size

To address how antibiotics affect the structure of gonococcal biofilms, we focused on early biofilms, namely (micro)colonies. Gonococci were allowed to assemble colonies within flow chambers for one hour and subsequently treated with antibiotics. Constant supply of nutrients and antibiotics was ensured by applying continuous flow. Prior to data acquisition, bacteria were stained with Syto 9 to detect the position of individual cells and to determine the cell volume (Fig. 1a). Moreover, propidium iodide (PI) enabled us to detect dead cells. Three-dimensional positions of single cells (Fig. 1b) were detected as described in the Methods section. This protocol allows analysing effects of antibiotic treatment on preassembled gonococcal colonies at single cell resolution providing spatial information.

**Fig. 1.**
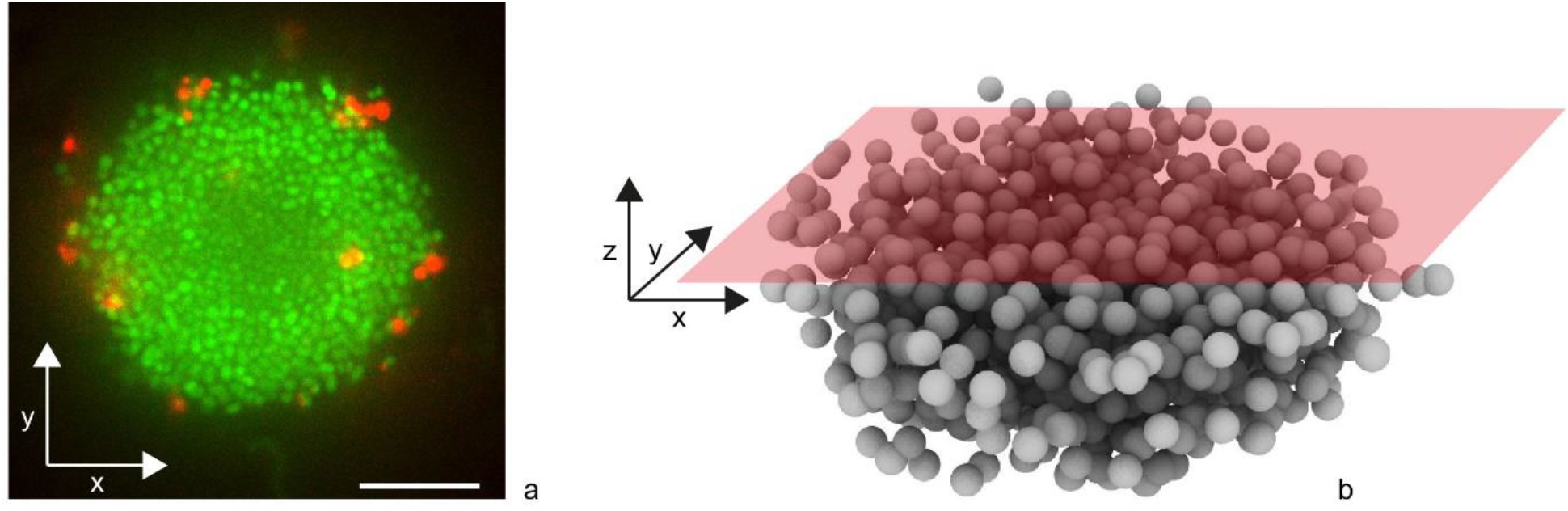
Determination of cell positions in three dimensions within spherical gonococcal colonies. a) Typical slice through Syto 9 (green)- and PI (red)-stained gonococcal colony. Scale bar: 10 µm. b) Confocal reconstruction of single cell positions.

We investigated the effect of the translation inhibitor azithromycin and cell wall synthesis inhibitor ceftriaxone, on the structure of colonies. After 5 h exposure to azithromycin or ceftriaxone at 100x their minimal inhibitory concentrations (MICs), the local structure changed considerably (Fig. 2a). Colonies had swollen and holes became prominent. To quantify these structural changes, we measured the radial distribution functions *g(r)*. For each bacterium, we determined the probability *g(r)* of finding a bacterium at a distance *r* (Fig. 2b). As previously shown [15, 16], gonococcal colonies show liquid-like local order, i.e. *g(r)* shows distinct maxima and minima at short but not at long distances *r* as seen in Fig. 2c for untreated and azithromycin-treated colonies. The position of the first maximum is the mean nearest neighbor distance *r*_*0*_. By fitting *g(r)* with an analytical expression for colloidal systems [27], we determined *r*_*0*_ (Fig. 2c, d). In the presence of azithromycin, the nearest neighbor distance *r*_*0*_ was strongly increased compared to the control (Fig. 2d), i.e. the entire colony had swollen. Under ceftriaxone treatment, we found no clear maxima in *g(r)* (Fig. 2c) indicating that liquid-like order was lost and the colony structure would be better characterized as gas-like. Independently, we measured the volumes of individual cells and found that for both antibiotics the volumes had increased by a factor of ∼ 2.5 compared to the control cells (Fig. 2e, Fig. S2). When the antibiotics were applied for a shorter period of time, we observed qualitatively similar effects on structure albeit less pronounced (Fig. S1, S2).

**Fig. 2.**
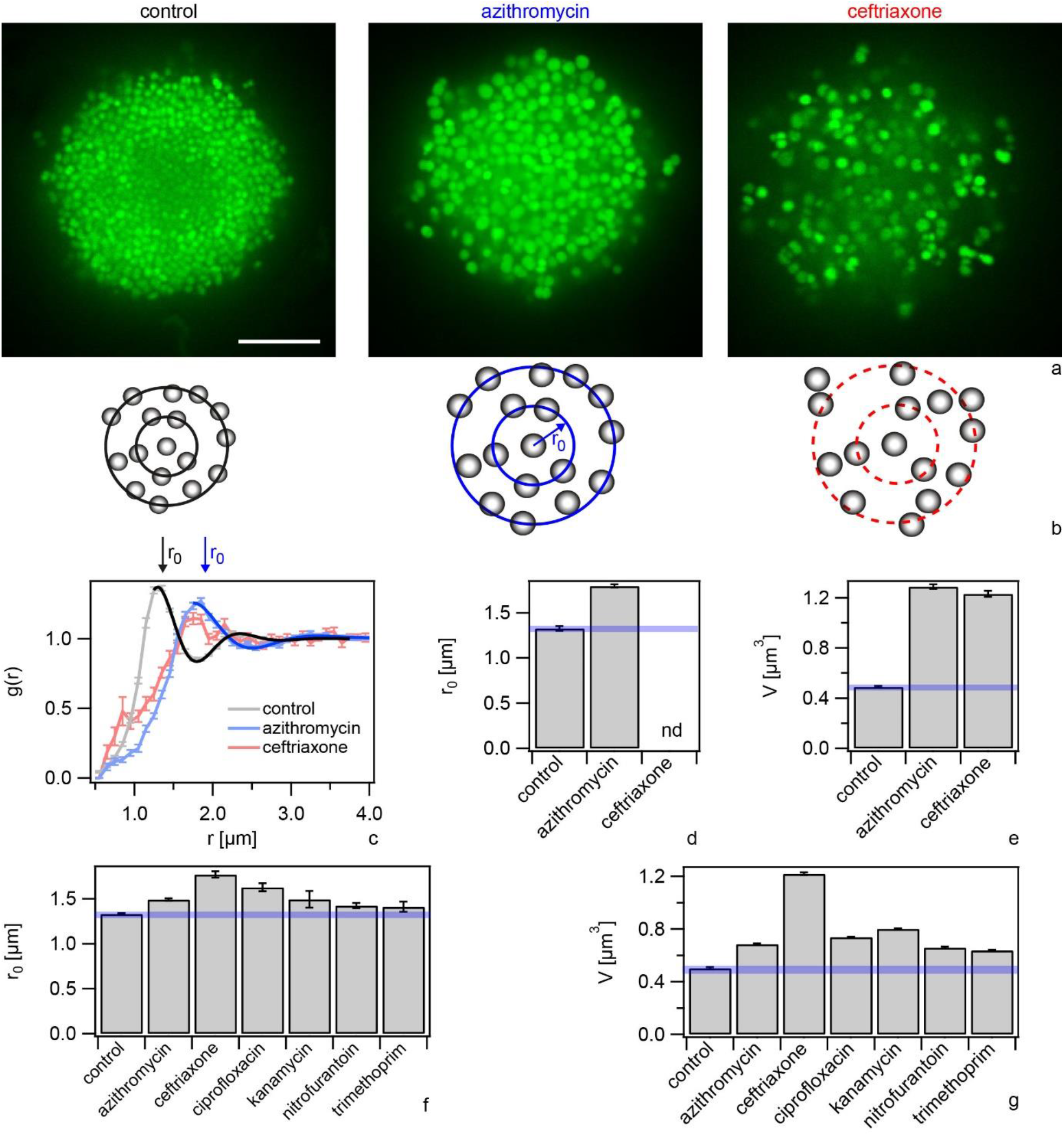
Antibiotics affect colony structure. Bacteria (wt*, Ng150) were inoculated into flow chambers and colonies were allowed to assemble for 1 h. Subsequently, they were treated with antibiotics at a-e) 100x MIC for 5 h, or f-g) 1x MIC for 3 h. a) Typical confocal slices. Scale bar: 10 µm. b) Scheme of local order. Left: Untreated cells show liquid-like order. The probability to find a nearest neighbor is highest at *r*_*0*_. Middle: azithromycin-treated cells show liquid-like order with larger *r*_*0*_. Right: ceftriaxone-treated cells show no local order. c) Radial distribution function *g(r)*. Error bars: standard error of the mean, N > 24 colonies. Full-line: Fit to eq. 1. d) Nearest neighbor distances *r*_*0*_ determined from g(r) (Fig. S1). Error bars: errors from fit to eq. 1. e) Mean cell volume *V*. Error bars: errors from Gaussian fits to distributions (Fig. S2), N > 20000 cells. f) Nearest neighbor distances *r*_*0*_ for 1x MIC determined from g(r) (Fig. S1). N > 24 colonies. Error bars: errors from fit to eq. 1. g) Mean cell volume *V* for 1x MIC. Error bars: errors from Gaussian fits to distributions (Fig. S2), N > 20000 cells. Blue transparent bars: uncertainties associated with mean values of controls.

To assess whether colony swelling was a general response of gonococci to antibiotics, we characterized the effects of antibiotics with different targets including translation (kanamycin), replication (ciprofloxacin), and folic acid metabolism (trimethoprim) as well as a producer of radicals (nitrofurantoin) on the structure of the colonies. These antibiotics were applied for 3 h at their respective MICs. All treatments increased the fractions of dead cells relative to the control (Fig. S3), however, most cells were still alive. Under these conditions, all colonies showed liquid-like local structure (Fig. S1), but the mean distance to the nearest neighbor *r*_*0*_ was shifted to higher values for all antibiotics tested (Fig. 2f). In agreement with an increased nearest neighbor distance, the mean cell volumes were significantly increased for all antibiotics tested (Fig. 2g, Fig. S4).

In summary, treatment with antibiotics with a large variety of targets increases the nearest neighbor distance in liquid-like colonies as well as the cell volumes.

### Antibiotic treatment affects the within-colony motility

Driven by T4P retraction, gonococci show motility within colonies (Fig. 3a, b, Movie S1). Here, we found that motility increased strongly under azithromycin treatment (Fig. 3, Movie S2). Single cell motility was characterized by tracking the positions of individual cells within a confocal plane. Fig. 3b shows a typical example of rapid movement of three cells at a time scale of seconds. We quantified motility by calculating the effective diffusion constant *D* by ⟨*x*(*t* = 1*s*)^2^⟩ = 4*D* · *t* from trajectories of single cells as shown in Fig. 3c. The effective diffusion constant tends to decrease with increasing distance from the edge of the colony (Fig. 3a, c, d) in agreement with previous reports [15, 28, 29]. Importantly, after 5 h of azithromycin treatment at 100x MIC, the effective diffusion constant was increased by a factor of ∼ 10 at the edge (Fig. 3e) and by a factor of ∼ 25 at 5 µm inside the colony (Fig. 3f).

**Fig. 3.**
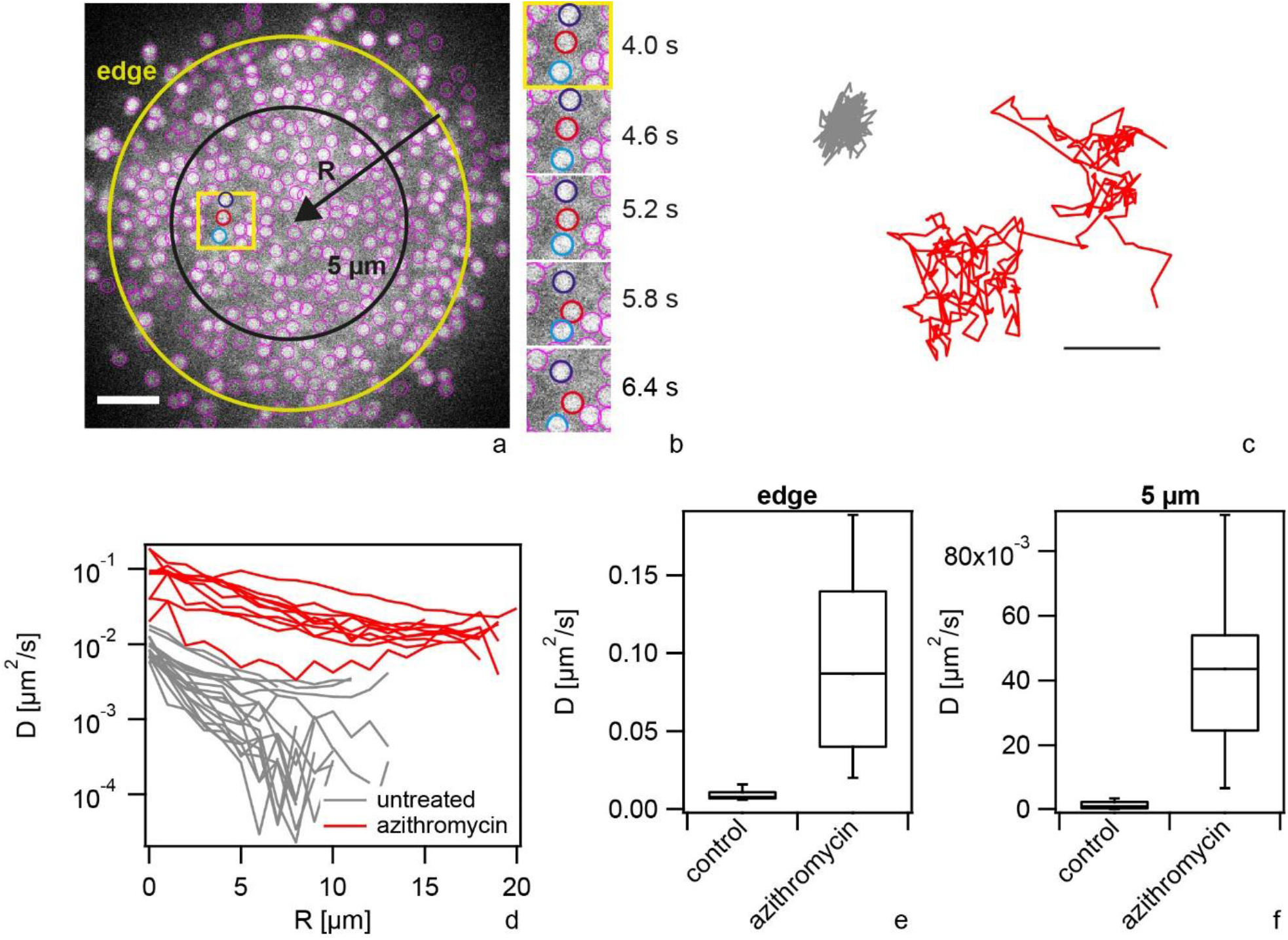
Within-colony motility increases during azithromycin treatment. wt* gonococci (Ng150) were inoculated into flow chambers and colonies were allowed to assemble for 1 h. Subsequently, they were treated with azithromycin (100x MIC) for 5 h. a) Typical confocal plane of wt* colony with Syto 9 staining. Purple circles: cells detected by tracking algorithm. Scale bar: 5 µm. b) Time lapse within region of interested depicted by yellow box in a). c) Typical traces of untreated (grey) and treated (red) single bacteria over 25 s. Scale bar: 0.5 µm. d) Effective diffusion constant as a function of penetration depth for untreated (grey) and treated cells (red), respectively. Effective diffusion constant *D* at e) the edge of the colony and f) within the colony at *R = 5* µm. Box plot: median, 25/75 percentile, whiskers: 10/90 percentile. N = 10 - 18.

We assessed whether treatment with lower concentrations or other antibiotics also affected the motility of gonococci within colonies (Fig. S5). After 3 h of azithromycin treatment at 1x MIC, the effective diffusion constant was increased by a factor of ∼ 3. Interestingly, ceftriaxone treatment significantly reduced the effective diffusion constant compared to the control. Ciprofloxacin strongly enhanced motility whereas nitrofurantoin showed no effect. For kanamycin and trimethoprim, the effects were different at the edge and at the center of the colony (Fig. S5). Taken together, various antibiotics enhance or inhibit within-colony motility.

### Azithromycin and ceftriaxone affect T4P-mediated attraction between neighboring cells

We hypothesized that reduced or enhanced T4P-T4P interactions govern the motility of cells within gonococcal colonies during antibiotic treatment. To assess this hypothesis, we characterized cell-to-cell interactions using a double laser trap. Again, we focused on azithromycin and ceftriaxone which show converse effects on motility. The laser traps were positioned at a distance of 2.8 µm and a single monococcus was trapped in each of the traps. When T4P of both cells attach to each other and at least one of them retracts, both cell bodies are pulled towards each other (Fig. 4a). Eventually, the bond ruptures and both cell bodies move rapidly back towards the centers of the traps. During several events, the cell bodies did not move back the full distance to the centers of the traps but instead paused or started retracting again (Fig. 4b). In this case, most likely multiple T4P-T4P bonds have formed simultaneously, and we detect only the shortest bond.

**Fig. 4.**
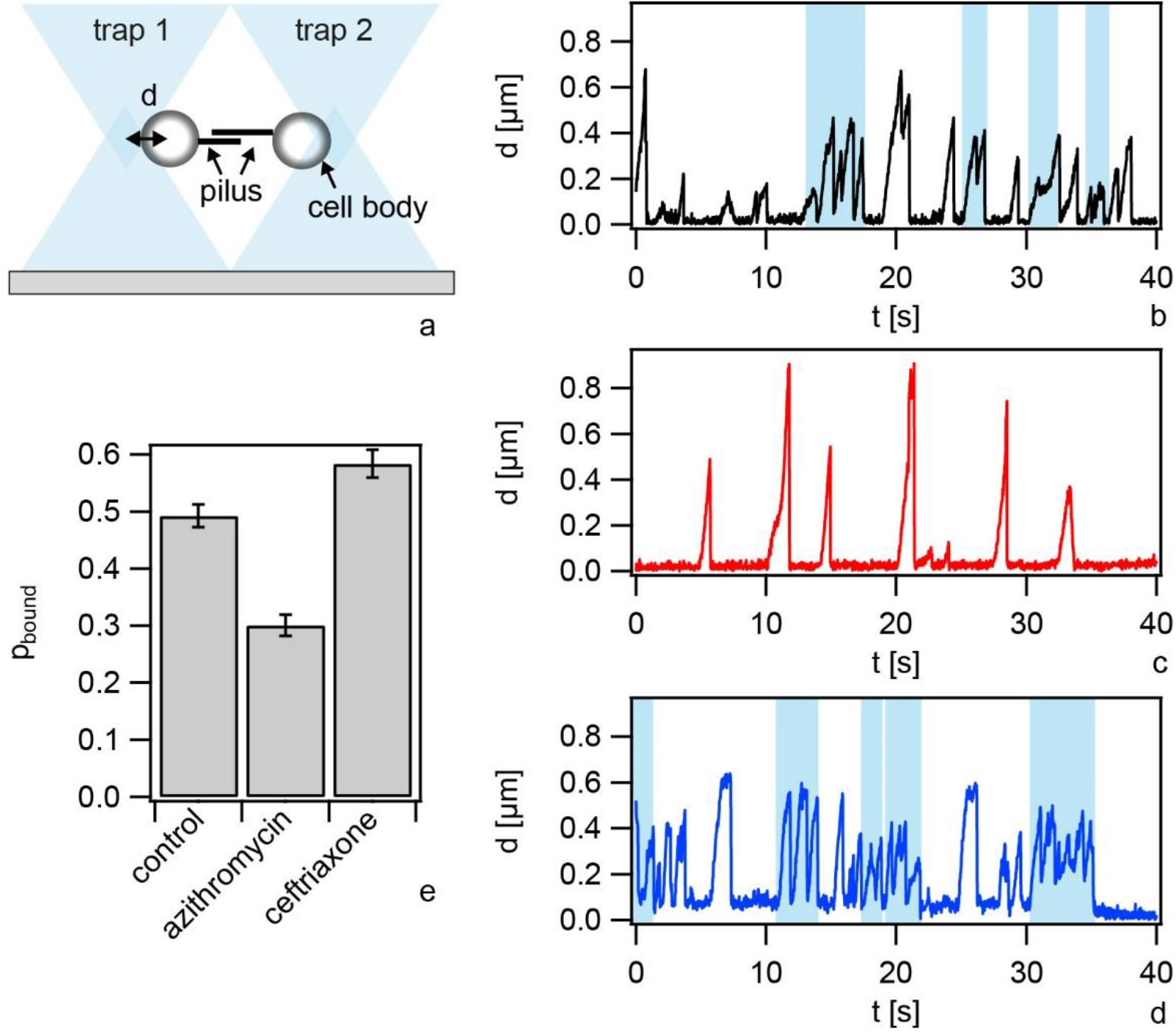
T4P-T4P interactions are reduced under azithromycin treatment and enhanced under ceftriaxone treatment. a) Sketch of the setup. A single bacterium is trapped in each of the traps. As T4P from different bacteria bind and retract, the cell bodies are deflected by the distances *d* from the centers of the traps. When the T4P-T4P bond ruptures, the cell bodies move rapidly back to *d = 0*. Typical dynamics of deflections b) without treatment c) with 6.4 µg / ml azithromycin for (2 - 3) h, and d) with 0.4 µg / ml ceftriaxone for (2 - 3) h. Blue shaded boxes: multiple T4P pairs exist simultaneously (bundling). e) Probability that cells are bound via T4P-T4P bonds. N = (29-38) pairs, error bars: bootstrapping.

We compared cells that were incubated for 1 h in medium and subsequently for 2 h in medium containing either azithromycin or ceftriaxone at 100x MIC with control cells that were incubated in medium for 3 h. We found less motor activity for azithromycin-treated pairs compared to the control pairs (Fig. S6) with the individual retraction events being well separated (Fig. 4b, c). By contrast, untreated pairs had a strong tendency towards multiple T4P-T4P bonds. Ceftriaxone treatment increased the frequency of T4P retractions and multiple T4P-T4P bond formation (Fig. 4d, Fig. S6).

Attractive interactions between pairs of cells were quantified by measuring the probability that two cells are bound to each other via T4P *p*_*bound*_. This probability was determined for each pair of bacteria by normalizing the total amount of time during which the bacteria are deflected from the trap centers by the total time elapsed between the first and the last retraction event. The mean probability to be bound was considerably lower for azithromycin-treated pairs with 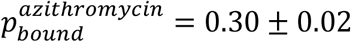 compared to the control pairs with 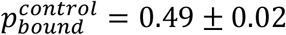 (Fig. 4e) and higher for ceftriaxone-treated pairs with 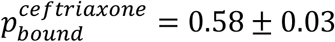. We note that our protocol is different from the previously used protocol where cells were directly harvested from agar plates explaining the difference in *p*_*bound*_[16] in the control samples.

We conclude that azithromycin treatment inhibits and ceftriaxone enhances T4P-mediated attractions between neighboring cells.

### Within-colony motility correlates with survivability

Having observed that antibiotic treatment strongly influences cell motility, we asked - vice versa - how cell motility affected the efficacy of antibiotic treatment. Recently, it was shown that gonococcal aggregation by means of T4P protects bacteria from ceftriaxone treatment [30]. Therefore, we wondered whether colony dynamics impacts on survivability and used bacterial strains that form less fluid colonies as revealed by characterizing the relaxation dynamics of colonies. Fluidity of gonococcal colonies can be affected by the motor activity of the T4P or by the binding strength between T4P [16, 17]. In particular, colony fluidity is strongly reduced in a strain expressing genes encoding for non-functional *pilT*_*WB*_ T4P retraction ATPases in addition to the functional ones [16]. Here, we verified that lower fluidity at the colony scale correlates with lower motility at the level of single cells. The effective diffusion constants of untreated cells were significantly lower in the *pilT*_*WB2*_ strain compared to the wt* strain (Fig. S7). To make sure that antibiotics-related effects are not merely caused by PilT functionality, we employed a second strain that reduces cell motility within colonies by a different mechanism. We used a strain lacking pilin phosphoform modification by deletion of *pptA* [31] resulting in reduction of colony fluidity [17]. Again, we found that single cell motility is reduced in strain *ΔpptA* compared to the wt* (Fig. S7).

First, we investigated the effect of cell motility on cell death in the absence of antibiotic treatment. The fraction of dead cells was determined by dividing the number of dead cells as detected by PI staining by the total number of cells detected by Syto 9 (Fig. 5a). Under benign conditions, strains *pilT*_*WB2*_ and *ΔpptA* showed a significantly higher fraction of dead cells compared to the wt*, suggesting that reduced motility was deleterious.

**Fig. 5.**
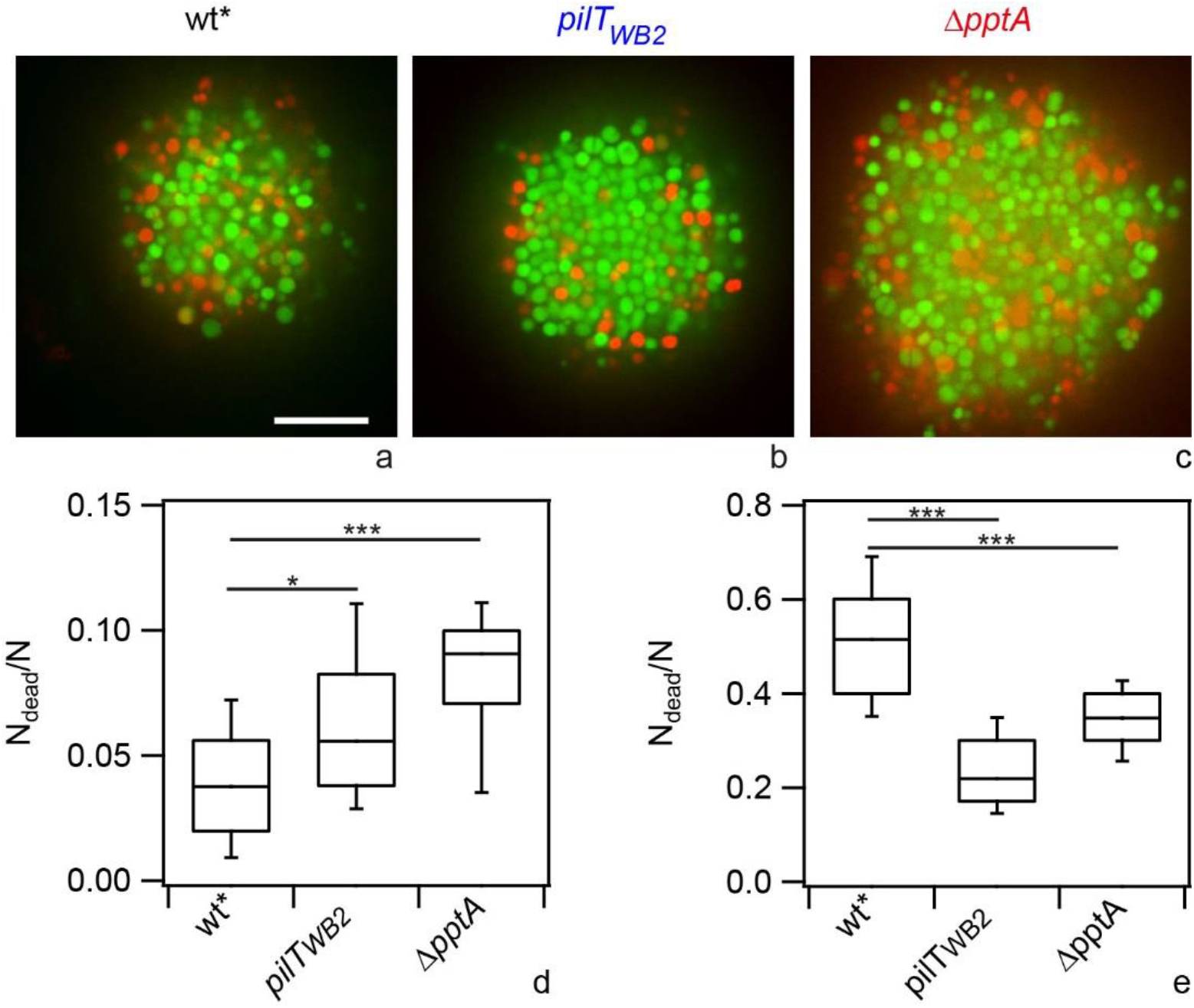
Fraction of dead cells in colonies with different fluidities. Bacteria were inoculated into flow chambers and colonies were allowed to assemble for 1 h. Subsequently, they were treated with 0.4 µg / ml ceftriaxone for 5 h. Typical confocal slices through ceftriaxone-treated colonies formed by a) wt* (Ng150), b) *pilT*_*WB2*_ (Ng176), and c) *ΔpptA* (Ng142). Green: Syto 9, red: PI. Scale bar: 10 µm. Fraction of dead cells d) untreated and e) with ceftriaxone. N = 32-45 colonies. All statistical comparisons were made by two-sample KS-test: *P 0.05; ***P 0.001.

Subsequently, we addressed killing by ceftriaxone treatment. After 5 h of treatment at 100x MIC, a fraction of f_dead_ = 52 % of the wt* cells within the colony were dead. Within the colonies formed by motility-inhibited cells, the fractions of dead cells were considerably lower with f_dead_ = 23 % for *pilT*_*WB2*_ and f_dead_ = 35 % for *ΔpptA* (Fig. 5e). At the edge of the colonies the fractions of dead cells of wt* and *ΔpptA* bacteria were comparable (Fig. S8), suggesting that bacteria are equally susceptible to antibiotic treatment when they are not surrounded by other bacteria. With increasing penetration depth *R, f*_*dead*_ decreased much more steeply for *ΔpptA* colonies compared to wt* colonies. This indicates that the less motile *ΔpptA* cells are better protected within the colony compared to the more motile wt* cells. At the edge of *pilT*_*WB2*_ colonies, the fraction of dead cells was lower compared to the wt* colonies. This suggests that even in the absence of neighboring cells, the *pilT*_*WB2*_ bacteria are less susceptible to ceftriaxone (Fig. S8a). Importantly, when the fractions of dead cells were normalized to the values at the edges of the colonies, *f*_*dead*_ showed a very steep decrease for both motility-inhibited colonies, *pilT*_*WB2*_and *ΔpptA* (Fig. S8b), showing that cell death was inhibited by colony formation. For very large depths *R > 15 µm*, the *f*_*dead*_ became comparable between all three strains.

Taken together we found that T4P mutant strains that reduce colony fluidity and within-colony motility show reduced survivability under benign conditions but strongly enhanced survivability under antibiotic treatment.

### Sequential treatment with azithromycin and ceftriaxone shows synergistic killing

So far, we found that treatment with various antibiotics including azithromycin enhances motility of bacteria within colonies. Also, our data strongly suggest that enhanced motility makes the bacteria more susceptible to ceftriaxone treatment. Therefore, we asked whether the killing efficiency of the drug combination azithromycin and ceftriaxone is additive. To address this question, we designed different treatment protocols all using 100x MIC with a total expose time of 3 h azithromycin and 3 h ceftriaxone.

First, we treated colonies with pure azithromycin for 3 h, and subsequently with pure ceftriaxone for another 3 h. After a total treatment for 6 h, the fraction of dead cells was determined (Fig. 6a). We found that the mean fraction of dead cells was 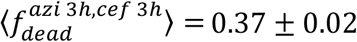. To find out whether the effect was additive, we measured 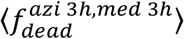 after 3 h of azithromycin treatment and 3 h of incubation in medium. Moreover, we determined 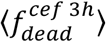 after 3 h of ceftriaxone exposure. Then we determined 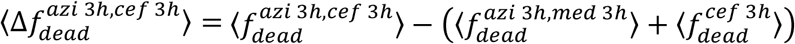. This is a measure of the fraction of dead cells arising from the interplay between both antibiotics. The first term on the right side describes the mean fraction of dead cells after combined treatment while the second term describes the sum of the mean fractions of dead cells after individual treatments. Interestingly, 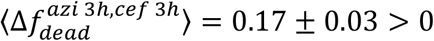 (Fig. 6d) indicates the sequential treatment with azithromycin and ceftriaxone is synergistic. Next, we inverted the order of the sequential treatment and found 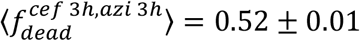 and 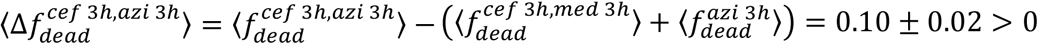 (Fig. 6b, d). Importantly, sequential treatment with both drugs shows synergistic effects regardless of the order of application. The total fraction of dead cells was higher when cells were treated first with ceftriaxone and then with azithromycin. However, the interpretation of this result is hampered by the fact that detection of cell death is delayed after ceftriaxone treatment (Fig. 6a, b). The fraction of dead cells determined directly after 3 h treatment was three-fold lower compared to the fraction determined after 3 h treatment and 3 h growth in medium. Therefore, we compared the degree of synergism plotted in Fig. 6d showing 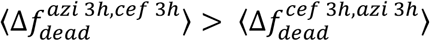.

**Fig. 6.**
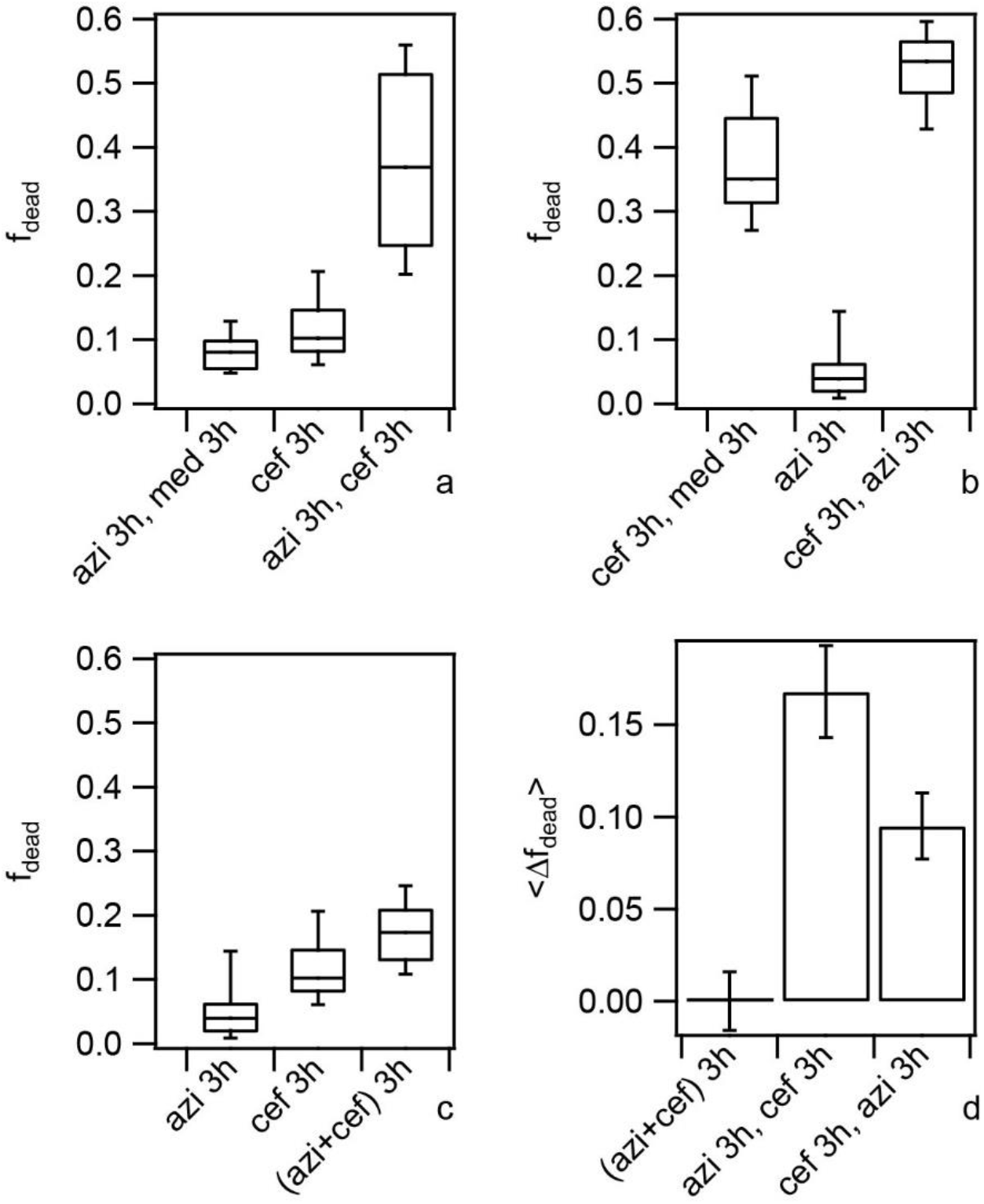
Sequence of drug application affects fraction of dead cells. Wt* bacteria (Ng150) were inoculated into flow chambers and colonies were allowed to assemble for 1 h. Subsequently, they were treated at 100x MIC with azithromycin or ceftriaxone, respectively, as indicated in the figure labels. a) - c) Fractions of dead cells *f*_*dead*_. Figure labels explain the protocol. For example, *azi 3h, med 3h* indicates that colonies were treated with azithromycin for 3 h, subsequently incubated with medium for another 3 h, and *f*_*dead*_ was determined after 6 h in total; *cef 3h* indicates that colonies were treated with ceftriaxone for 3 h and *f*_*dead*_ was determined after 3 h. Box plots: median, 25/75 percentile, whiskers: 10/90 percentile. N = (30 - 61) colonies. d) Difference <Δ*f*_*dead*_> between <*f*_*dead*_> of combined treatment and independent treatments was calculated as describe in the Results. Error bars: standard error of the mean.

Finally, we treated colonies simultaneously with azithromycin and ceftriaxone. We found 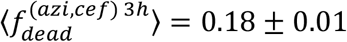 and 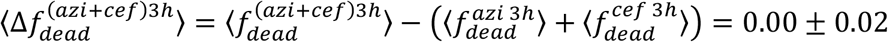 (Fig. 6c, d). This result indicates that the effects of azithromycin and ceftriaxone are additive during simultaneous treatment.

We conclude that sequential treatment with azithromycin and ceftriaxone (and vice versa) shows synergistic effects.

## Discussion

We present a single-cell level study of antibiotic effects on gonococcal colonies and focus on the interplay between cell motility and antibiotic treatment. Our method opens up the possibility to characterize novel parameters as compared to classical assays such as the detection of the fraction of viable cells by colony counting. First, we can characterize spatial effects within colonies. Second, cell dynamics and biophysical properties are accessible. Third, we characterize the instantaneous state of bacteria within colonies. This is particularly relevant for studying drug combinations because we probe bacteria at the moment of drug application. Fourth, we exclusively probe gonococci within colonies. Gonococci continuously produce unpiliated cells by pilin phase- and antigenic variation [32]. These cells are sorted out of the colonies [19, 33] and exist as planktonic cells. In plating assays, a mixture between colonie-associated and planktonic cells is probed. Despite these important benefits our method shows limitations compared to viable cell counting. One limitation associated with confocal microscopy is the penetration depth of several tens of micrometers which precludes the analysis of thick biofilms. Moreover, the determination of the fraction of dead cells is based on a fluorescence assay that probes membrane permeability. It is conceivable, that cells are unable to recover from antibiotic treatment even though their membrane is still intact at the time point of image acquisition. In this case, we would correctly determine that the cell is still alive but it would not produce offspring. In a plating assay these cells would be counted as dead cells. Indeed, for ceftriaxone treatment, we found delayed killing (Fig. 6) and this delayed killing would reduce colony counts. For azithromycin treatment, little cell death was observed even after 5 h treatment at 100x MIC. It is possible that while most of these cells have intact membranes after 5 h, they will never escape from their stressed state and resume cell growth. As a consequence, cells with an intact cell membrane but incapable of dividing would lead to lower counts in a plating assay. Characterizing both the short-term cellular response investigated in this study and long-term viability will provide a more complete picture of how antibiotics affect colony-bound gonococci.

Many bacterial species employ T4P for cell motility and aggregation [12]. In *Neisseria* species, T4P-T4P mediated attractive forces control the dynamics of bacteria within colonies [15-17, 28]. Here, we show that treatment with the protein synthesis inhibitor azithromycin reduces T4P-dependent attraction and, conversely, ceftriaxone enhances the attraction. This result is consistent with the following picture (Fig. S11). A gonococcus continuously extends and retracts its pili. Extended pili bind to pili of multiple neighboring cells. When a pilus retracts, the cell moves. During movement force builds up, eventually leading to rupture of the pilus-pilus bond. Thus bacterial motility within colonies can be understood as a continuous tug-of-war between adjacent bacteria [12]. Azithromycin treatment reduces the probability that a pilus-pilus bond is formed or maintained. As a consequence, the restoring forces from neighboring bacteria are reduced and bacteria become more mobile. Ceftriaxone has the opposite effect of enhancing the probability that neighboring cells are bound via T4P thus reducing within-colony motility.

Increased single cell motility within colonies correlates with increased colony fluidity (or decreased viscosity) [15-17]. Fine-tuning T4P-T4P mediated attractive forces induces a transition between gas-like, liquid-like, and solid-like behavior of the colonies. For *N. meningiditis* fluidity was correlated with their success in colonizing narrow tubing [15]. In particular, it was shown that motile cells that form fluid-like colonies were more efficient in colonizing blood vessels than non-motile cells forming solid-like colonies. Here, we demonstrate that cell motility correlates with fitness in a different manner, namely by influencing survivability. Ceftriaxone treatment was less effective within motility-inhibited colonies compared to highly motile colonies. Different mechanisms could explain our result. First, motility could enhance the influx of macromolecules as was shown for *Bacillus subtilis* biofilms [20]. Under benign conditions, nutrient influx would be faster. Our experimental observation that survivability of motile cells is higher would agree with this hypothesis. In the presence of antibiotics, motility-inhibited colonies would slow down antibiotic penetration. Considering the fairly small size of our colonies, however, diffusion seems unlikely to be a limiting factor. Second, reproducing bacteria require space and bacteria residing at the center of the colony would be unable to grow unless the surrounding bacteria are motile enough to rearrange and create space. When motility is inhibited, central bacteria are likely to respond to growth inhibition by switching into the stationary state. In the growth-inhibited state, dead bacteria would accumulate explaining why the fraction of dead cells is high for motility-inhibited cells in the absence of antibiotics. In the presence of antibiotics, growth inhibition prior to the application of antibiotics is likely to confer tolerance and protect central bacteria from killing. Thus, we consider stress response due to limited space in the center of motility-inhibited colonies as likely explanation for reduced killing in these colonies. Taken together, our study together with the study of Bonazzi et al [15] demonstrates that within-colony motility confers strong fitness effects. High motility enhances survivability and the ability to colonize vessel-like confinements. However, it reduces survivability during antibiotic treatment.

Here we found that azithromycin treatment enhances cell motility and that increased cell motility makes colony-bound bacteria more susceptible to ceftriaxone treatment. This would suggest that treatment by a combination of both drugs was synergistic. Indeed, we found evidence for synergism with sequential treatment but not with simultaneous treatment. The synergistic effect was highest when colonies were first treated with azithromycin and subsequently with ceftriaxone. This result is consistent with azithromycin increasing cell motility, thus boosting the killing efficiency of ceftriaxone. Further studies will be required to find out why the reversed order of drug application also shows synergism.

For planktonic rod-shaped bacteria it is well-known that the cell morphology changes when bacteria are treated with certain antibiotics [34, 35]. When *Escherichia coli* is treated with β-lactams or fluoroquinolones, it strongly increases its cell length [34, 36]. By contrast, treatment with protein synthesis inhibitors has only minor effects on cell volume [34, 37]. In biofilm-dwelling rod-shaped *V. cholerae*, antibiotics inhibiting protein synthesis were shown to enlarge the cell volume considerably [11]. While protein synthesis (and consequentially cell division) was inhibited, metabolism was still active causing the cell volume to increase. Curiously, we are unaware of any study investigating the effects of antibiotics on the cell volume of spherical bacteria (cocci). In line with filamentation of rod-shaped cells [34, 35], we found that the cell volume of gonococci increases strongly (2.5 fold) when treated with the β-lactam antibiotics ceftriaxone at its MIC. When we applied high doses of the protein synthesis inhibitor azithromycin for extended periods of time, the cell volume approached the volume of ceftriaxone-treated bacteria, in agreement with the reports on *V. cholerae* [11]. Additionally, we found that also the folic acid inhibitor trimethoprim and the radical producing nitrofurantoin increased the cell volume. The fact that antibiotics from five different classes with four different targets induce an increase in cell volume when applied at the MIC suggests that cell size increase might be a result of general stress response. Inhibition of cell division is observed in response to various stresses [38] and could explain why cells grow larger. An increase in cell volume has been implicated with increased antibiotic tolerance enabling cells to extend their survival time during antibiotic treatment [39]. We conclude that spherically shaped gonococci increase their cell volume in response to treatment with different classes of antibiotics.

In conclusion, we demonstrate that antibiotics do not only affect biofilm structure but also its dynamics and identify T4P-mediated cell-to-cell attractions as the underlying mechanism. Since many biofilm-forming species form T4P [12], this finding likely has implications beyond the *Neisseria* field. Decreased cell-to-cell attractions enhanced the efficiency of antibiotic killing within colonies. In future studies it will be interesting to find out whether this holds true also for bridging attractions other than T4P.

## Materials and Methods

### Growth conditions

Gonococcal base agar was made from 10 g/l BactoTM agar (BD Biosciences, Bedford, MA), 5 g/l NaCl (Roth, Darmstadt, Germany), 4 g/l K_2_HPO_4_ (Roth), 1 g/l KH_2_PO_4_ (Roth), 15 g/l BactoTM Proteose Peptone No. 3 (BD Biosciences), 0.5 g/l soluble starch (Sigma-Aldrich, St. Louis, MO), and supplemented with 1% IsoVitaleX (IVX): 1 g/l D-glucose (Roth), 0.1 g/l L-glutamine (Roth), 0.289 g/l L-cysteine-HCL x H_2_O (Roth), 1 mg/l thiamine pyrophosphate (Sigma-Aldrich), 0.2 mg/l Fe(NO_3_)_3_ (Sigma-Aldrich), 0.03 mg/l thiamine HCl (Roth), 0.13 mg/l 4-aminobenzoic acid (Sigma-Aldrich), 2.5 mg/l β-nicotinamide adenine dinucleotide (Roth), and 0.1 mg/l vitamin B12 (Sigma-Aldrich). GC medium is identical to the base agar composition but lacks agar and starch.

### Bacterial colony formation and flow chambers

Bacterial colonies were grown in ibidi sticky-Slides I^0.8^ Luer flow chambers with a glass bottom at a constant nutrient flow of 3 ml/h by using a peristaltic pump. Bacteria were harvested from overnight plates and resuspended in GC medium to an optical density at 600 nm (OD600) of 0.05, 300 µl of the cell suspension was inoculated into the chambers. The flow chambers were incubated at 37 °C. Antibiotics were added by changing the medium at the inlet.

### Measurement of minimal inhibitory concentrations

The MICs were measured in 48 well plates (Greiner Bio-One) by measuring the OD600 using a plate reader (Infinite M200, Tecan). Each well was inoculated with 5 × 10^5^ CFU/ml and antibiotics were added. The well plate was incubated for 24 h at 37 °C and 5% CO_2._ The lowest concentration that did not lead to a change in OD600 over 24 h was determined to be the MIC. To avoid effects of colony formation, MICs were determined for strain *ΔpilE* (Ng196) that does not generate T4P and therefore colony formation is severely suppressed for this strain [19]. For comparison, we measured the MICs of the colony-forming wt* strain for azithromycin and ceftriaxone. As expected, they were higher compared to the MICs measured with the aggregation-inhibited strain.

### Confocal microscopy

Prior to image acquisition, colonies were stained using 500 µl of GC medium containing 3 µmol/l Syto 9 (Thermo Fisher Scientific) and 4.5 µmol/l propidium iodide (PI) (Thermo Fisher Scientific). Images were acquired using an inverted microscope (Ti-E, Nikon) equipped with a spinning disc confocal unit (CSU-X1, Yokogawa) and a 100x, 1.49 NA, oil immersion objective lens. The excitation wave lengths were 488 and 561 nm.

### Determination of bacterial positions in three dimensions

Single cell positions were determined in three dimensions using a method described in [16]. In short, confocal image stacks were filtered using a spatial bandpass filter. In the resulting image, local maxima were determined yielding the bacterial coordinates with pixel accuracy. Subpixel positions were determined by calculating the centroid of spherical masks around the local maxima.

### Determination of radial distribution function

The radial distribution function *g(r)* is defined so that *N*/*V g*(*r*)*r* ^2^*dr* is the probability that the center of a bacterium will be found in the range *dr* at a distance *r* from the center of another bacterium, where *N*/*V* is the number density of bacteria. To calculate *g(r)*, we calculated the distribution of distances between cells *n(r)* by determining the distances between all pairs of cells and sorting them into bins with width dr. The distribution *n(r)* was then smoothed using a moving average filter to generate the distribution *N(r). N(r)* is an approximation of the distribution of distances of a system with the same density as for *n(r)* but with random distances between the particles. The radial distribution function was then calculated via the relation *g*(*r*) = *n*(*r*)/*N*(*r*).

As described previously [16], the shapes of *g(r)* show good qualitative agreement with the radial distribution functions found in colloidal systems with Lennard-Jones-like interactions. We used the formula proposed by Matteoli and Mansoori [27]

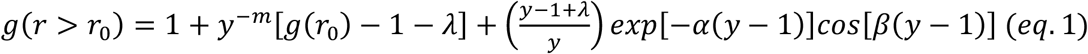

where *r* _0_ is the nearest neighbor distance, *y* = *r* /*r* _0_ and *m, λ, α*, and *β* are adjustable parameters.

### Determination of fraction of dead cells

The fraction of dead cells was computed by dividing the number of dead cells by the total number of cells found within a colony. The number of dead cells within a colony was determined by counting the cells found in the PI channel. The total number of cells was measured by determining the sum of cells found in the PI and Syto 9 channel. In addition to the total fraction of dead cells per colony, the fraction of dead cells in dependence of the distance to the colony contour was measured. A binary mask of the colony was generated using an intensity threshold. The contour of the mask was extracted representing the colony contour. The closest point of the contour was determined for each cell and the distance was calculated. The cells were sorted into bins according to their distance to the contour and the fraction was determined as described above.

### Determination of cell volume

The volume detection was based on an algorithm presented by [40]. A Laplacian of Gaussian filter was used to highlight the cell boundary. Masks representing the cellular objects were generated by filling the boundary and Watershedding was applied to separate falsely connected objects and diplococci. Then, the volume of the masks was determined yielding the volume of the cells.

To verify that the observed cell volume increase was not merely an effect of chromosome rearrangements, we imaged single cells in phase contrast. Overnight cultures were resuspended in GC medium plus Isovitalex and incubated for at least 1 h in liquid medium (37°C, 5% CO_2_) with a starting optical density of OD_600_= 0.05. Subsequently, antibiotics were added at the appropriate concentration. After 2 h of treatment, bacteria were diluted 1:10 in phosphate buffered saline (PBS), put onto coverslips sealed with VALEP and subsequently imaged using phase contrast microscopy. The size of a bacterium was determined by analysing the intensity profiles of individual cocci using *ImageJ*. In agreement with the fluorescence data, cells treated with azithromycin were larger than control cells (Fig. S4). We note that the total volume estimated from this experiment is expected to be different from the volume measured using confocal microscopy because it depends on the exact definition of the edge of the cell. The diffraction pattern of the phase contrast image introduces an uncertainty of the absolute value. However, we are only interested in relative changes in radii under antibiotic treatment.

### Time-resolved single cell tracking in two dimensions

Trajectories of single cells in time were obtained using the Fiji (Fiji is just ImageJ) Plugin TrackMate in the following manner: Spinning Disc Confocal Microscopy time series at mid-height were loaded into Fiji, regularized using the Plugin StackReg (Rigid body regularization), then divided by a Gaussian smoothed version with smoothing radius 9 px (corresponding to ∼ 1 cell diameter). In TrackMate, the threshold intensity and apparent cell diameter *d* was set to adequately capture the cells, but not any noise (we generally set *d* to 1 mum for normal sized cells, and *d* to 1.3 mum for enlarged cells). Further, the maximal displacement in the algorithm allowed for cells between two time frames was set to 0.4 mum. The resulting tracks were then saved as a CSV table.

### Measurement of effective diffusion constant

From the two dimensional tracking data obtained in Fiji, the effective diffusion coefficients were determined as following: From the trajectories, the cell positions and velocities, as well as the center of mass of all cells, were corrected for global colony translation and rotation in Matlab by subtracting the average translational and angular velocity from the velocity of each cell. From here, *D*(*r*) is obtained as the mean-square displacement at *t* = 1*s* divided by 4. of all tracked cells with track length larger than 3 time steps (frame rate 50 Hz).

Static errors in the determination of particle positions (due to a noisy intensity signal) were obtained and corrected by calculating the velocity auto-correlation function *VACF*(*t, r*). The latter showed clear decorrelation of single cell dynamics for *t* > *dt*, and we thus assumed that *VACF*(*dt, r*) = − *E*/2*dt*^2^, with *E* equal the error of the MSD which we then substracted.

### Characterization of T4P-mediated cell-to-cell attraction

In order to determine the interactions on the single-cell level we used a dual laser trap setup that was slightly modified from [16]. The trapping laser (20I-BL-106C, 1064 nm, 5.4 W; Spectra Physics, Santa Clara, CA) was directed into a water-immersion objective (Nikon Plan Apochromate VC 60× NA 1.20; Nikon). Manipulation of the trap position was done with a two-axis acousto-optical deflector (DTD-274HD6 Collinear Deflector; IntraAction, Bellwood, IL). The incoming signal for the AOD was generated using a signal generator (SDG 2042 X Function/Arbitrary Waveform Generator, Siglent, Germany) and reinforced by a synthesizer. The distance between the centers of the two traps was adjusted to be 2.84 μm, corresponding to 1.6 MHz in our setup. The beam was focused onto the sample plane via an 60x-objective (Plan Apochromate VC 60x water immersion, N.A. 1.20, W.D. 0.27mm, Nikon, Japan). Bacterial dynamics was recorded with a high-speed camera (EoSens 3CL, Mikrotron, Germany) with a framerate of 50 frames per second. The position of each bacterium for every point in time was calculated using the Hough transformation and MATLAB, described in detail in [16]. From the displacement tracks, durations and frequencies of retraction, elongation, pausing and bundling states were determined.

The samples were prepared by selection of single piliated colonies from overnight GC-plates and resuspended in GC medium plus Isovitalex. After 3h of incubation (37°C, 5% CO_2_) in liquid medium in a well with a starting optical density of OD_600_= 0.05, the bacteria were diluted 1:10 in GC medium plus Isovitalex and ascorbic acid (500 mM). Prior the addition of antibiotics, bacteria were grown at least for 1 h in liquid media liquid in a well. Then antibiotics were added with the appropriate concentration. After 2 h of treatment, bacteria were prepared like the control strain.

## Acknowledgements

We thank Knut Drescher and Anton Welker for suggestions about single cell detection, Tobias Bollenbach for critical reading of the manuscript, and Paul Higgins and the Maier lab for helpful discussions. This work was supported by the Deutsche Forschungsgemeinschaft through grant MA3898 and the Center for Molecular Medicine Cologne.

## Notes

### Competing Interest Statement

The authors have declared no competing interest.

