## Supplementary figures & Tables for "Antibiotics modulate attractive interactions in bacterial colonies affecting survivability under combined treatment"

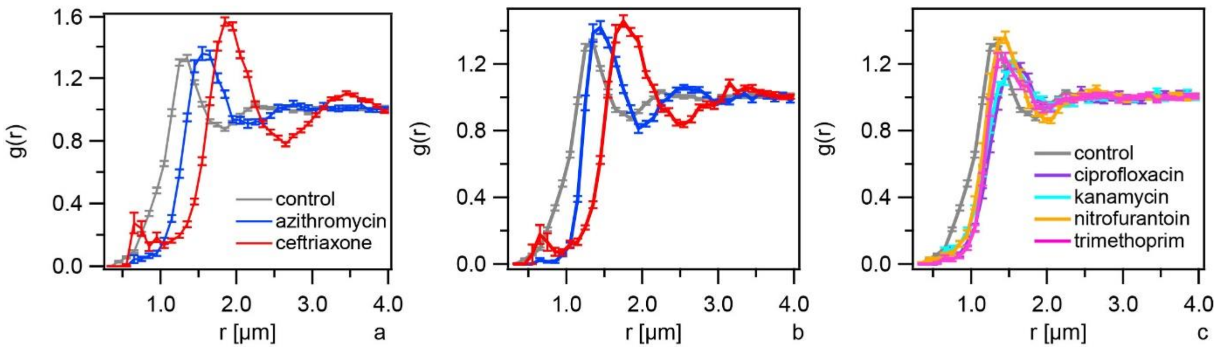


Fig. S1 Radial distribution functions. Bacteria (wt*, Ng150) were inoculated into flow chambers and colonies were allowed to assemble for 1 h. Subsequently, they were treated with antibiotics for 3 h at a) 100x MIC, or b, c) 1x MIC. The values for $r_0$ shown in Fig. 2d, f are obtained from fits to eq. 1 to these distributions.

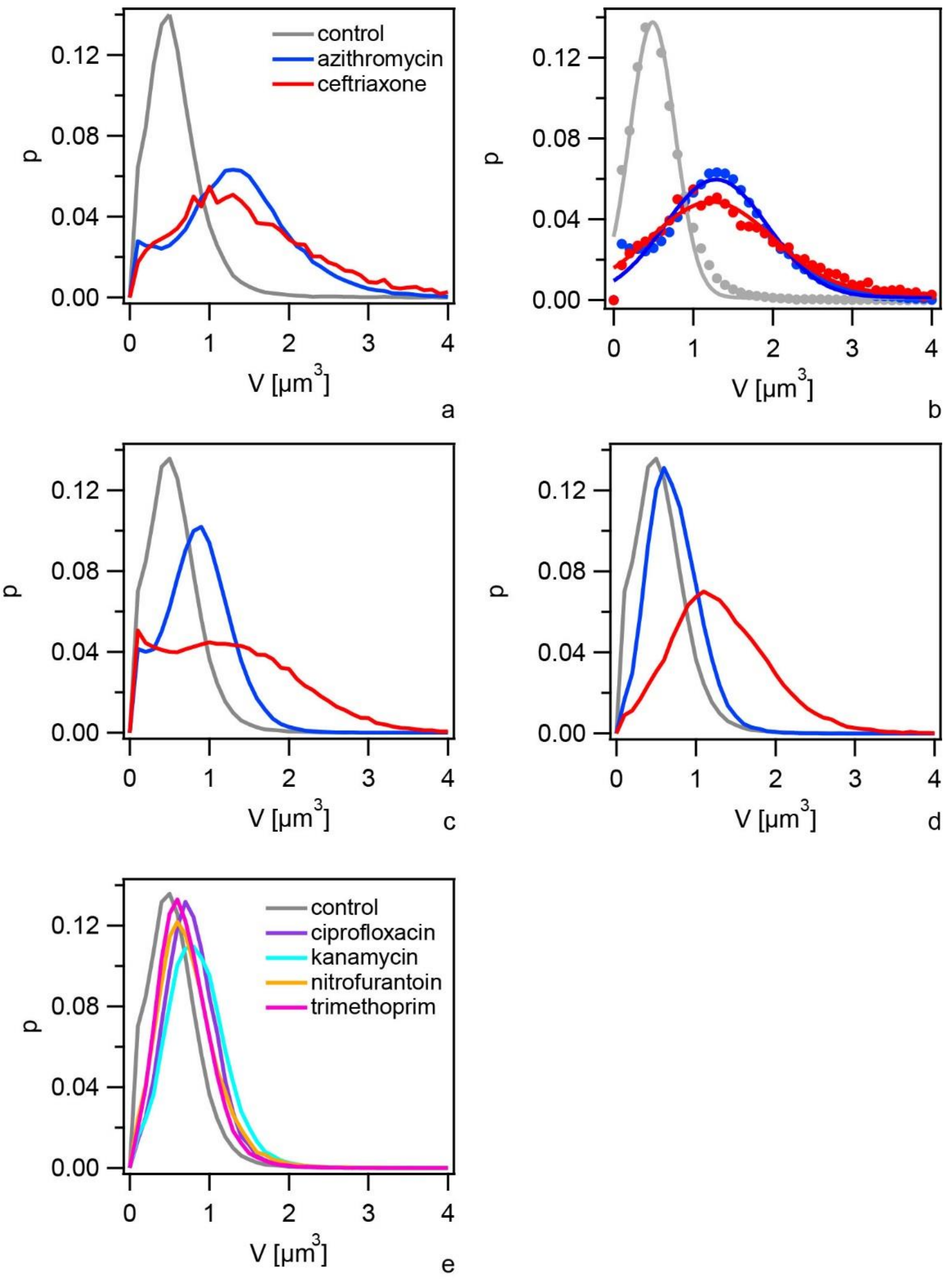


Fig. S2 Distributions of cell volumes. Bacteria (wt*, Ng150) were inoculated into flow chambers and colonies were allowed to assemble for 1 h. Subsequently, they were treated with antibiotics for a, b) 5 h at 100x MIC, c) 3 h at 100x MIC, d - e) 3 h at 1x MIC. The mean values shown in Fig. 2e, g are obtained from Gaussian fits to these distributions. b) Markers: data points, full lines: fits to Gaussian function.

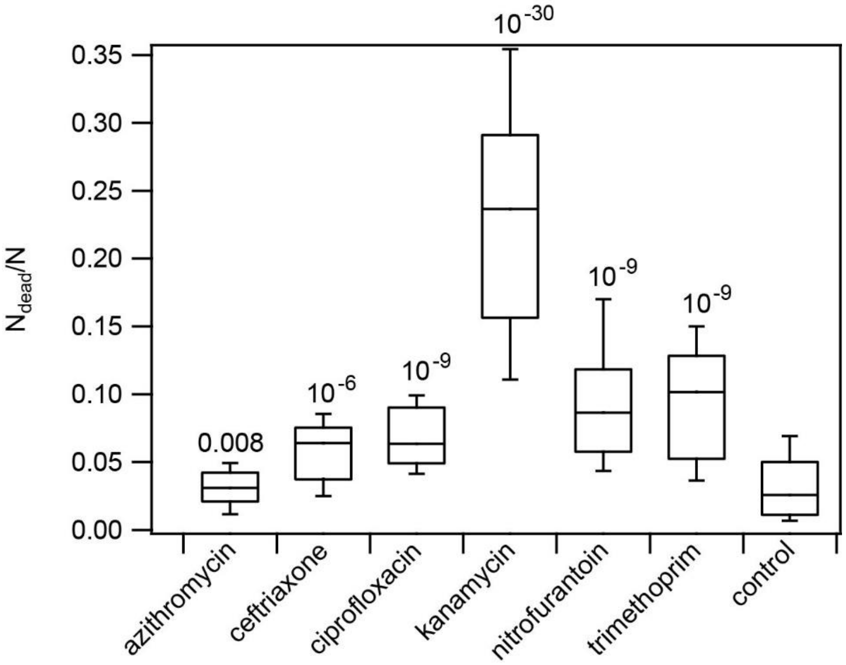


Fig. S3 Fractions of dead cells $f_{dead} = N_{dead}/N$ after 3 h of treatment at 1x MIC. Numbers: p-values from two sample KS-test against the control.

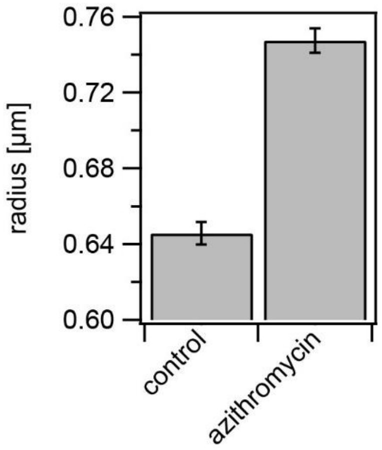


Fig. S4 Control for cell radius increase under azithromycin treatment. Wt* cells (Ng150) were incubated for 1 h in liquid medium and subsequently treated with azithromycin at 100x MIC for 2 h. Radii of individual cells were determined using phase contrast microscopy. Mean ± standard error of radii of N = 75 cells. $p < 10^{-16}$ (KS test).

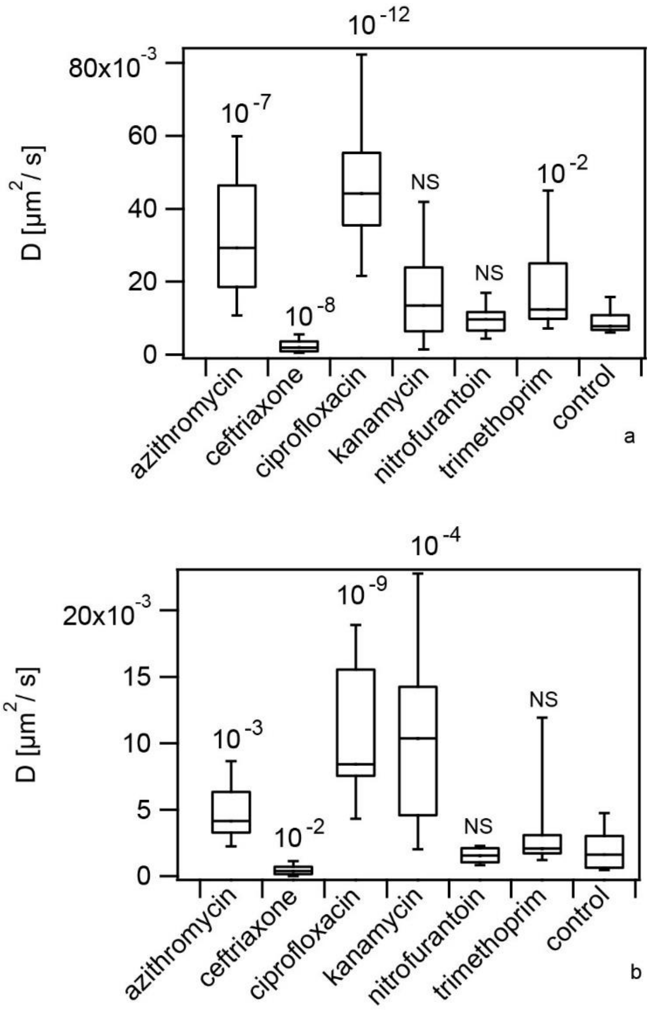


Fig. S5 Antibiotic treatment strongly affects bacterial motility within colonies. wt* gonococci (Ng150) were inoculated into flow chambers and colonies were allowed to assemble for 1 h. Subsequently, they were treated with different antibiotics at their respective MICs for 3 h (0.064 µg / ml azithromycin, 0.004 µg / ml ceftriaxone, 0.002 µg / ml ciprofloxacin, 20 µg / ml kanamycin, 0.48 µg / ml nitrofurantoin, 32 µg / ml trimethoprim). Effective diffusion constant was measured at a) the edge and b) at R = 5 µm into the colony. N = 12 - 20 colonies. Numbers are p-values from two sample KS-test against the respective controls.

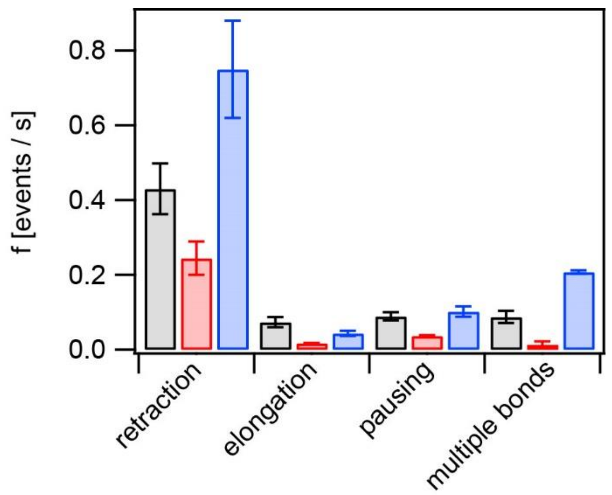


Fig. S6 Frequencies of T4P retraction, elongation, pausing, and multiple T4P-T4P bonds. Grey: control, red: with 6.4 µg / ml azithromycin for (2 - 3) h, blue: with 0.4 µg / ml ceftriaxone for (2 - 3) h. Error bars: bootstrapping.

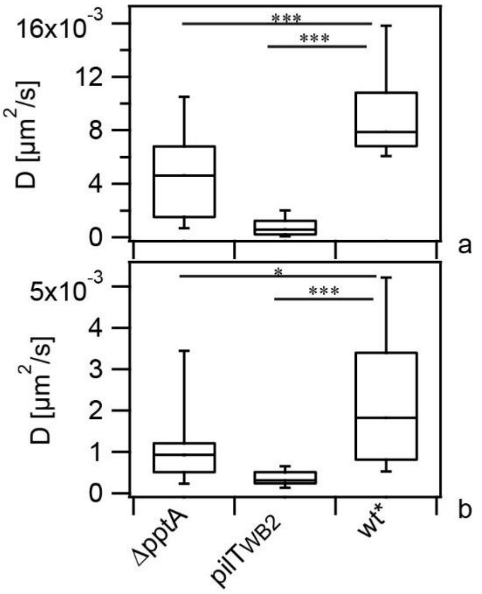


Fig. S7 Mean effective diffusion constant *D* of untreated cells at a) the edge of the colony and b) within the colony at *R* = *5* µm after 6 h of growth for strains *ΔpptA* (Ng142), $pilT_{WB2}$ (Ng176), and wt* (Ng150). Box: 25/75 percentile, whiskers: 10/90 percentile. N = 13 - 20. All statistical comparisons were made by two-sample KS-test: *P 0.05; ***P 0.001.

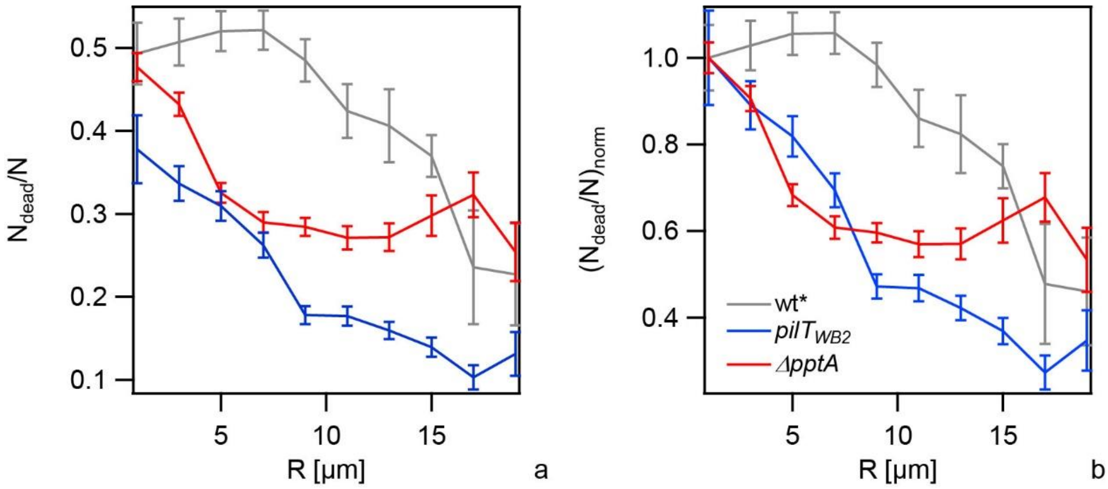


Fig. S8 Fraction of dead cells $f_{dead} = N_{dead}/N$ in colonies as a function of distance from the edge of the colony at R = 0. Bacteria were inoculated into flow chambers and colonies were allowed to assemble for 1 h. Subsequently, they were treated with with 0.4 µg / ml ceftriaxone for 5 h. a) Fractions of dead cells. b) Fractions of dead cells normalized to fraction at the edge. Grey: wt* (Ng150), blue: *pilT<sub>WB2</sub>* (Ng176), red: *ΔpptA* (Ng142). Error bars: standard error of the mean. N = (9 - 46) colonies per data point.

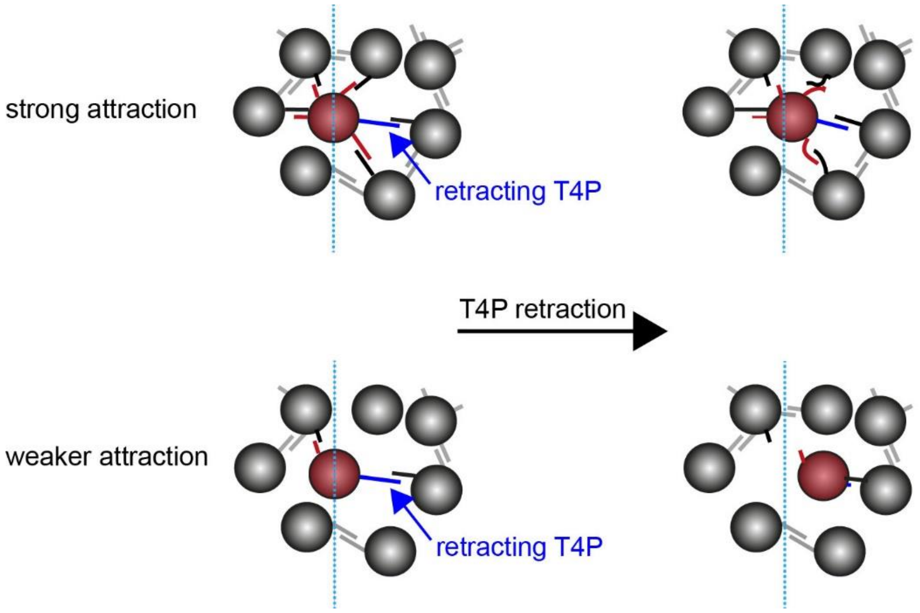


Fig. S9 Tug-of-war model for motility within colony. In the absence of azithromycin, a bacterium within the colony simultaneously forms multiple bonds with adjacent bacteria (top). When a T4P retracts, movement of the cell body is hindered by T4P-T4P bonds at the opposite side of the retracting pilus. In the presence of azithromycin, the probability that a T4P-T4P bond is formed is reduced (bottom). Thus the probability that a retracting T4P has opposing T4P-T4P bonds is lower. As a consequence, the bacterium is more motile.

### Supplementary Tables

| Strain | Relevant genotype | Source/Reference |
|---|---|---|
| *wt** (Ng150) | *G4::aac* | [17] |
| $pilT_{WB2}$ (Ng176) | *iga::*$P_{pilE}$ *pilTWB ermC*<br>*G4::aac* | [16] |
| *ΔpptA* (Ng142) | *pptA::kan*<br>*G4::aac* | [17] |
| *ΔpilE* (Ng196) | *pilE::cat*<br>*G4::aac* | This study, [16] |

Table S1 Bacterial strains used in this study.

| **antibiotic** | **MIC [µg / ml]** *ΔpilE* | **MIC [µg / ml]** wt* |
|---|---|---|
| **azithromycin** | 0.064 | 0.128 |
| **ceftriaxone** | 0.004 | 0.008 |
| **ciprofloxacin** | 0.48 | |
| **kanamycin** | 32 | |
| **nitrofurantoin** | 0.002 | |
| **trimethoprim** | 20 | |

Table S2 Minimal inhibitory concentrations (MICs) of antibiotics. MICs were determined from bacteria that cannot form colonies (*ΔpilE*, Ng196) and from colony-forming *wt** (Ng150) by testing for the ability to grow overnight.
